# Statistical learning of frequent distractor locations in visual search involves regional signal suppression in early visual cortex

**DOI:** 10.1101/2021.04.16.440127

**Authors:** Bei Zhang, Ralph Weidner, Fredrik Allenmark, Sabine Bertleff, Gereon R. Fink, Zhuanghua Shi, Hermann J. Müller

## Abstract

Observers can learn the locations where salient distractors appear frequently to reduce potential interference – an effect attributed to better suppression of distractors at frequent locations. But how distractor suppression is implemented in the visual cortex and frontoparietal attention networks remains unclear. We used fMRI and a regional distractor-location learning paradigm (Sauter et al. 2018, 2020) with two types of distractors defined in either the same (orientation) or a different (colour) dimension to the target to investigate this issue. fMRI results showed that BOLD signals in early visual cortex were significantly reduced for distractors (as well as targets) occurring at the frequent versus rare locations, mirroring behavioural patterns. This reduction was more robust with same-dimension distractors. Crucially, behavioural interference was correlated with distractor-evoked visual activity only for same- (but not different-) dimension distractors. Moreover, with different- (but not same-) dimension distractors, a colour-processing area within the fusiform gyrus was activated more when a colour distractor was present versus absent and with a distractor occurring at a rare versus frequent location. These results support statistical learning of frequent distractor locations involving regional suppression in the early visual cortex and point to differential neural mechanisms of distractor handling with different-versus same-dimension distractors.

## Introduction

In everyday life and experimental scenarios, such as, the additional-singleton paradigm (Theeuwes 1992), attention is often distracted or ‘captured’ by salient but goal-irrelevant stimuli (Folk and Remington 1998; Hickey et al. 2006; Forster and Lavie 2008). However, with repeated exposure and practice (Kelley and Yantis 2009; Zehetleitner et al. 2012), distractor interference can be reduced via attentional control (Bacon and Egeth 1994; Leber and Egeth 2006; Müller et al. 2009; Gaspelin et al. 2017).

Moreover, observers can learn not only to prioritize locations for attentional selection where task-relevant targets are regularly encountered (Shaw and Shaw 1977; Geng and Behrmann 2005), but also to deprioritize locations where salient but irrelevant distractors frequently appear (Goschy et al. 2014; Leber et al. 2016; Ferrante et al. 2018; Sauter et al. 2018; Wang and Theeuwes 2018). Typically, in the latter studies, a salient distractor occurs with a higher likelihood at one, ‘frequent’ display location/subregion relative to the remaining, ‘rare’ locations/subregions. The consistent finding is that, over time, search becomes less impacted by distractors that appear at frequent than at rare locations. This effect is attributable primarily to a proactive suppression of frequent distractor locations: oculomotor capture is less likely when distractors occur at frequent (vs. rare) locations (Di Caro et al. 2019; Wang et al. 2019a; Sauter et al. 2020); and for frequent locations, an anticipatory suppression-related event-related component (P_D_) is observed (Wang et al. 2019b). However, how suppression of likely distractor locations is implemented is influenced by how distractors are defined relative to the target (Sauter et al. 2018; Allenmark et al. 2019; Failing et al. 2019; Zhang et al. 2019; Liesefeld and Müller 2020): if target and distractor are defined in the same dimension (e.g., target and distractor are both orientation-defined), suppression appears to work at a supra-dimensional level of ‘attentional-priority’ computation, impacting both distractor and target signals – as compared to a level of dimension-specific ‘feature-contrast’ computation when they are defined in a different dimension (orientation-defined target, colour-defined distractor), in which case suppression typically impacts only distractor signals.

While a consensus is emerging as to the loci of learnt distractor-location suppression within the architecture of search guidance, how suppression is neurally implemented remains largely unclear. It is well-established that the frontoparietal network, including the inferior/superior parietal lobe (IPL/SPL), is involved in attentional control of distractor interference (de Fockert et al. 2004; Krueger et al. 2007), and top-down control can instigate preparatory activity to minimize capture by expected distractors (Serences et al. 2004; Ruff and Driver 2006; Munneke et al. 2011). For instance, presenting trial-by-trial precues indicating the likely target side as well as, on critical trials, the appearance of a distractor in the opposite hemifield, Ruff and Driver (2006) observed enhanced occipital-cortex activation in the hemisphere contralateral to the upcoming distractor during the cue period, and this was associated with reduced search costs later on. Yet, concerning top-down effects on distractor coding in early visual cortex, the evidence is mixed. For instance, Bertleff et al. (2016) found precuing of the target region to diminish distractor interference through increased activity in medial parietal regions involved in controlling spatial attention, rather than by down-modulating distractor signals in early visual cortex. In contrast, manipulating the overall likelihood with which a distractor could occur anywhere in the display, Won et al. (2020) reported distractor signalling in visual cortex to be diminished when distractors occurred architecture, along with reduced distractor interference.

Thus, using functional magnetic resonance imaging (fMRI) in Sauter et al.’s (2018, 2020) regional distractor-location learning paradigm, the current study aimed to examine whether visual-cortex signals at learnt distractor locations would be down-modulated to reduce distractor interference and what specific role the frontoparietal attention networks play in distractor handling. In particular, given the dissociative learning effects between distractors defined in the same versus a different dimension to the target, we examined differences in neural mechanisms mediating distractor-location learning between the two distractor types.

## Materials and Methods

### Participants

Thirty-two volunteers (mean age: 27.47 years; age range: 20-45 years; 18 female) were recruited, twenty-four at the Forschungszentrum Jülich and eight at the LMU Munich. Functional MRI data from six participants were excluded from the MRI analysis due to data quality (e.g., distortion) issues and/or head movements. Based on the effect size of significant preparatory visual activation of distractor suppression in Serences et al. (2004), for a power of 0.80 and an alpha of 0.05 (G*Power analysis) (Erdfelder et al. 1996), the required sample size would have been 24. However, to attain enough power and take into account potential drop-outs, 32 subjects were recruited. All participants were right-handed and reported normal or corrected-to-normal vision, including normal colour vision, and none had been diagnosed with a neurological or psychiatric disorder. Participants received 15 Euro per hour for their service. The study protocol was approved by the ethics committees of the German Society of Psychology (DGPs) and, respectively, the Psychology Department of the LMU Munich. Written informed consent was obtained from all participants before the experiment.

### Apparatus

In preparation for the fMRI experiment, participants received behavioural training outside the scanner to become familiarized with the task. The training was conducted in a sound-reduced and moderately lit test chamber. Stimuli were presented on a 24-inch Samsung SyncMaster 2233 (Samsung Electronics Co., Ltd., Seoul, South Korea) screen at a 1280 × 1024 pixels screen resolution and a 120-Hz refresh rate. Stimuli were generated by Psychophysics Toolbox Version 3 (PTB-3) (Brainard 1997) based on MATLAB R2019 (The MathWorks® Inc). Participants viewed the monitor from a distance of 60 cm (eye to screen), and distance and fixation position were controlled by a forehead-and-chin rest and an EyeLink 1000 eye-tracker device. In the experiment proper (in the scanner), stimuli were presented on a 30-inch LCD screen mounted behind the scanner 245 cm away from the head coil. The stimulus settings and the parameters for MRI data acquisition were the same at the Forschungszentrum Jülich and the LMU Munich.

### Visual Search Task

#### Stimuli

The stimuli used were essentially the same as in Sauter et al. (2018, 2020). The visual search displays consisted of twenty-nine turquoise (CIE [Yxy]: 29.6, 0.23, 0.37, measured on an equivalent display outside the scanner) upright or inverted ‘i’ shaped bars (0.10° of visual angle wide, 0.50° high; see search display in Figure 1A). One bar was positioned in the centre of the screen; the other bars were arranged on three imaginary concentric circles (around the centre) with radii of 1.25°, 2.50°, and 3.75° of visual angle containing 4, 8, and 14 items, respectively. The target was an item defined by a unique orientation difference compared to the vertically oriented nontarget items: it was tilted 30° to either the left or the right, with tilt direction randomized across trials. On a fraction of trials, one of the nontarget items, the singleton distractor (referred to as ‘distractor’ hereafter) was defined by either a different colour (red; CIE [Yxy]: 29.7, 0.30, 0.27; the *different-dimension distractor*) or a different orientation (a 90°-tilted, i.e., horizontally oriented ‘i’, the *same-dimension distractor*) compared to all the other items. The target and the singleton distractor only appeared at one of the eight positions on the middle circle, and they never appeared at the same location or adjacent to each other. The nontarget items on the outer and inner rings served to equate local feature contrast amongst the various singleton positions. All search items were presented on a black screen background (3.58 *cd*/ *m*^2^).

**Figure 1.**
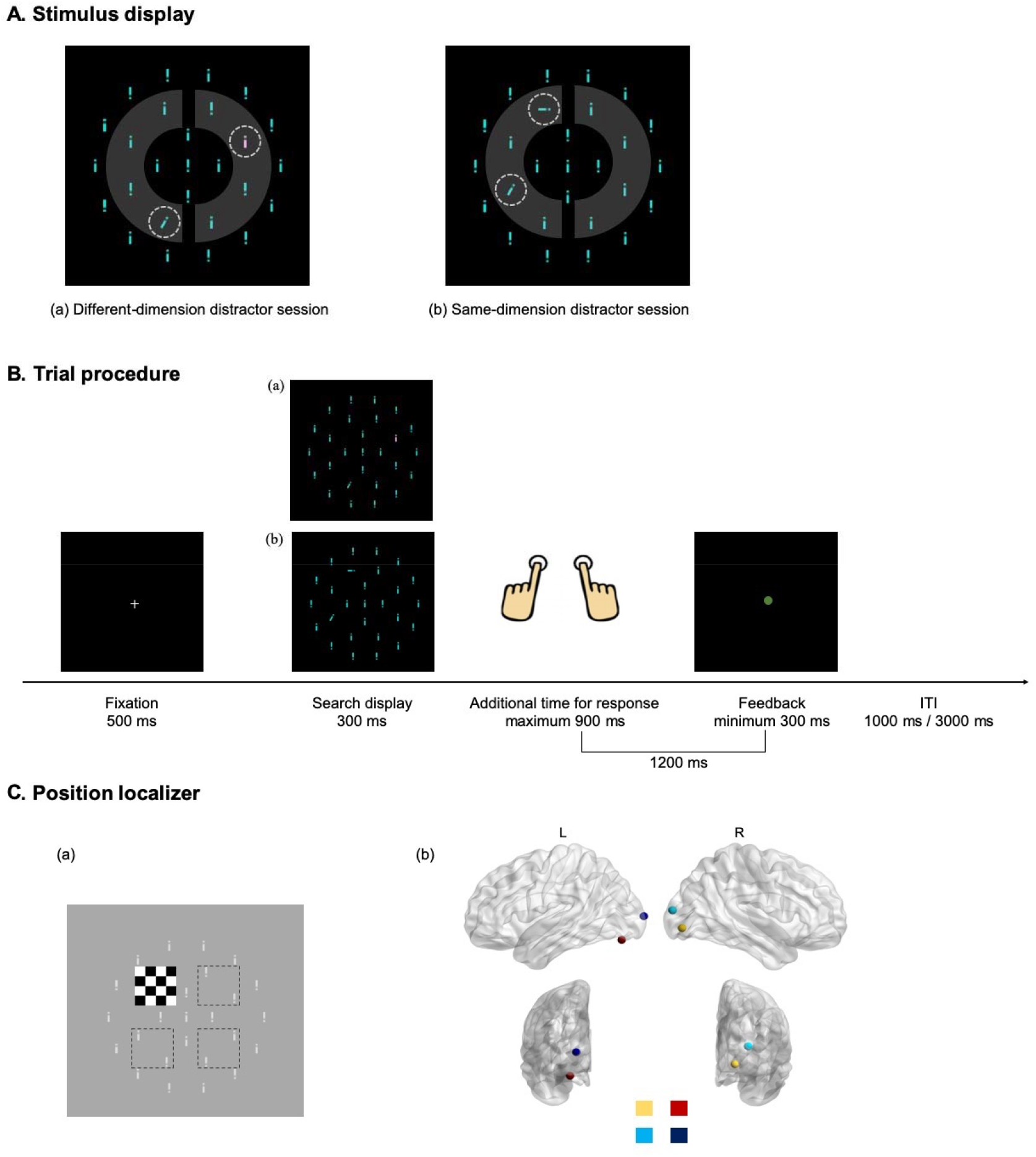
**A**. Example of a search display in (a) the different-dimension distractor session: the search target is the 30-°tilted item (here, outlined by a white dashed circle, bottom-left of the middle ring), and the distractor is a red colour singleton (outlined by a white dashed circle, top-right of the middle ring); (b) the same-dimension distractor session: the search target is again the 30°-tilted item (outlined by a white dashed circle, bottom-left of the middle ring), and the distractor is a horizontal orientation singleton (outlined by a white dashed circle, top-left of the middle ring). Grey-shaded areas indicate the eight potential target and distractor locations, and the left and right grey semicircles indicate the frequent and, respectively, rare distractor regions. Note that the dashed lines and grey areas are for illustration purposes only; they were not shown in the experiment. **B**. Example of the trial procedure, described in more detail in the text. **C**. (a) Example of a checkerboard stimulus (here, top-left) serving as positional localizer for possible target and distractor locations; note that the black dashed-line squares and the grey ‘i’ stimuli are depicted here only for illustration purposes, i.e., they were not presented in the experiment. (b) The four VOIs induced by the position localizers, coded by four colours, are projected onto a brain surface rendering.

Note that the physical bottom-up saliency of the two types of distractors was determined in a pilot study (with different participants) in which the colour and, respectively, the orientation distractor were presented as response-relevant targets; that is, in separate blocks, they were the only singleton item in the display, to which participants had to make an eye movement as fast as possible. Following Zehetleitner et al. (2013), we took the (saccadic) reaction time to indicate the physical saliency of a given distractor stimulus. Results revealed that a similar proportion of first saccades was directed to the red and the horizontal target, 92% and 90%, respectively. Latencies of the first saccade were somewhat shorter for the red compared to the horizontal target, 166 ms vs. 184 ms, *t*(8.06) = -2.93, *p =* .019, *d*_*z*_ *=* 1.69. Thus, taking the two measures together: if anything, the physical saliency of the red singleton was somewhat higher than that of the horizontal singleton.

#### Design

The two types of singleton distractor were introduced as a session factor in a within-group design: participants encountered only one type of distractor, either a different-dimension (i.e., colour) or a same-dimension (i.e., orientation) distractor, in either the first or the second experimental session (with order counterbalanced across participants). In each session, the singleton distractor was presented in 60% of trials, the remaining 40% being distractor-absent trials. If a distractor was present, it appeared with 80% probability in one half of the search display (i.e., at one of the four positions on the middle semicircle on either the left or the right side – henceforth referred to as the ‘frequent’ distractor region) and with 20% probability in the other half (the ‘rare’ distractor region) (see Figure 1A). In contrast to the distractor, the target appeared equally often in both regions, with an equal probability for all eight possible positions.

Note that for each participant, the region in which the distractor appeared frequently was reversed between two experimental sessions (e.g., if the left half was frequent in session 1, the right half was frequent in session 2), to rule out carry-over of learning effects between the two types of distractors. The assignments of the frequent distractor region (left vs. right semicircle) and the type of distractor (same dimension vs. different-dimension first) to the two sessions were counterbalanced across participants, thus avoiding possible confounds.

Further of note, the distractor type was manipulated as a within-subject factor in the present study: our participants had to learn the spatial distribution of one type of distractor first and then, after an unlearning phase, the opposite distribution with the other type of distractor. In previous studies (Sauter et al. 2018, 2019, 2020), we had used a between-subject design to avoid carry-over of acquired suppression strategies from one distractor type to the other. Based on finding dissociative target-location effects between same- and different-dimension distractors, we had proposed that statistical learning of distractor locations typically involves different levels of priority computation: the supra-dimensional priority map with same-dimension distractors (producing both a distractor- and a target-location effect) vs. a level specific to the distractor-defining dimension with different-dimension distractors (producing only a distractor-location effect). Despite possible carry-over effects potentially weakening dissociative effects between the two distractor types, for the present fMRI study, we opted for a within-participant design to examine statistical distractor-learning effects within the same brain. Also, note that with different-dimension distractors, both dimension- and priority-map-based suppression are in principle feasible. Thus, even if observers start with a priority-map-based strategy (as indicated by them displaying a target-location effect), most will revert to a dimension-based strategy (as indicated by observers losing the target-location, but not the distractor-location, effect) over extended task practice (Zhang et al. 2019).

#### Procedure

Each trial began with the presentation of a fixation cross in the middle of the screen for 500 ms, followed by the search display for a fixed duration of 300 ms (see Figure 1B). Participants were instructed to respond to the top vs. bottom position of the dot in the target ‘i’ by pressing the corresponding (left-/right-hand) response button (with stimulus-response assignment counterbalanced across participants) with two hands. Responses were to be made within 1200 ms of search display onset; otherwise, the trial was ‘timed out’. Following the response or time-out, feedback was provided in the shape of a coloured dot (0.4° of visual angle in diameter) presented in the screen centre: a green dot (RGB: 0, 255, 0) following a correct response and a red dot (RGB: 255, 0, 0) following an incorrect response or a time-out (i.e., too slow a response). A total time of 1200 ms was fixed for response and feedback: an additional maximum of 900 ms for response and a minimum of 300 ms for feedback (i.e., the feedback duration depended on the response time on a given trial). The intertrial interval (ITI) was 1000 ms or 3000 ms, randomly determined on each trial. Each experimental session consisted of 440 trials in total, subdivided into eight blocks of 55 trials. Five trials in each block only showed the search display without target and distractor which were treated as missing trials. Breaks of 6 s duration separated the blocks. Before the MRI scanning, participants performed three training blocks outside the scanner (with the same type of distractor as in the first experimental session) to practice the task (i.e., finding the target ‘i’ and responding to the dot position within it) and start learning the biased (80%/20%) spatial distractor distribution (to increase the power for determining the brain regions involved in statistical distractor location learning in the scanner). Besides, before practising the second session (also outside the scanner), participants completed four blocks (40 trials in each block) in which the singleton distractor was the same as in the first session but appeared equally often at two distractor regions (50%/50% distribution), to unlearn the spatial bias acquired for the first type of distractor. The number of unlearning trials was based on Ferrante et al. (2018), who found the distractor-location learning effect to be near-abolished within 144 ‘extinction’ trials.

In all experimental phases, participants were instructed to maintain fixation on the centre of the screen from the fixation cross’s appearance to the trial’s end. During practice (outside the scanner), compliance with this instruction was checked by monitoring participants’ eye movements using an eye-tracker device. In the scanner, eye movements could not be recorded, but participants reported that they had successfully maintained fixation in the vast majority of trials. Note also that making eye movements would actually have been counterproductive given the brief (300-ms) display duration.

### Position Localizer Task

To functionally identify the visual cortical representations corresponding to the various target and singleton distractor locations, a separate position localizer run was performed either before or after experimental session 1 (counterbalanced across participants). Participants were instructed to fixate the cross in the screen centre. They were then exposed to a contrast-reversing flickering checkerboard pattern that consisted of black and white mini-tiles (RGB: 0, 0, 0 and RGB: 255, 255, 255, respectively) flickering counter-phase at 8 Hz, with a height and width of 2°, which was presented successively in each quadrant of the visual field (see Figure 1C, left). Note that the localizer size covered two adjacent (target/distractor) locations in the search display. The localizer stimuli cycled through the four quadrants in clockwise direction, appearing at each location for 16 s with a 16 s break in-between for complete rounds, so that the localizer run took 4.27 min to complete.

### MRI Measurement and Analysis

#### Data acquisition

MRI data were acquired on a 3.0 T TRIO Prisma MRI (Siemens, Erlangen, Germany) whole-body MRI system equipped with a 64-channel head matrix coil. Each participant was fitted with cushions in the head coil to help stabilize the head position. Participants viewed the monitor via an adjustable mirror positioned on top of the head coil. Functional images were obtained using a blood oxygenation level-dependent (BOLD) contrast sensitive gradient-echo echo-planar sequence. A total of 1355 images were acquired in each experimental session and, respectively, 244 images in each positional localizer run; scanning parameters: TR = 1200 ms, TE = 30 ms, flip angle = 70 degree, FOV = 192×192 mm, voxels size = 2 × 2 × 3 mm, slices number = 36, slice thickness = 3 mm. Structural MRI images (T1-weighted) were acquired from the sagittal plane a using three-dimensional magnetization-prepared rapid gradient-echo (MP-RAGE) pulse sequence; scanning parameters: TR = 1780 ms, TE = 2.51 ms, flip angle = 8 degree, FOV = 256 × 256 mm, voxel size = 0.9 × 0.9 × 0.9 mm, slice thickness = 0.9 mm.

#### Preprocessing

Functional-imaging data were processed with SPM12 (r7771) (Wellcome Centre for Human Neuroimaging, London, United Kingdom; https://www.fil.ion.ucl.ac.uk/spm/software/spm12) based on MATLAB R2019a. Functional images acquired in each experimental session were corrected for interslice time differences for every participant first. Next, the functional images from the main experiment and those from the position localizer functional images were corrected for head movement by affine registration in a two-pass procedure realigning individual functional images to their mean image. Participants who exhibited translation head motion of more than 3 mm or rotations of more than 3 degrees were excluded from further analysis. Each participant’s mean image was then spatially normalized to a standard Montreal Neurological Institute (MNI) template using the ‘unified segmentation’ approach, and the resulting deformation field was applied to the individual functional images. The resulting images were smoothed with a 6-mm full width at half maximum (FWHM) Gaussian kernel to improve the signal-to-noise ratio and compensate for residual anatomical variations.

#### fMRI Analysis

Due to data quality issues (e.g., distortion) or large head movements during the visual search task, six out of the thirty-two participants were excluded from the functional MRI data analysis. To maximally use the available data, we included their good-quality behavioural and positional localizer data in the analysis.

##### Whole-brain analysis

The first-level (individual-participant) analysis involved applying a general linear model (GLM), with the following regressors for each distractor-type session. There were four primary regressors, one for each of the four basic experimental conditions of theoretical interest: two regressors for distractor-present trials, namely, singleton distractor in the frequent region and singleton distractor in the rare region; and two regressors for distractor-absent trials, namely, target in the frequent region and target in the rare region. Also, the two manual-response conditions (left button press, right button press) were included as regressors to avoid a high implicit baseline (Monti 2011), along with an extra regressor for unused trials (the first trial in each block and trials with incorrect/missing responses). The hemodynamic response related to neural activity in each of the above conditions was modelled by the canonical hemodynamic response function and its first derivative, which can capture the late negative dip of empirical BOLD responses (Henson et al. 2002). Finally, six head-movement parameters were considered as covariates in the model to reduce potential confounding effects (Lund et al. 2005). Based on the GLM, combining the same regressors across the two experimental sessions, we defined and calculated four contrast images at the first level to examine the effects of distractor interference (distractor present > distractor absent, and vice versa) and of distractor-location learning (distractor in the rare region > distractor in the frequent region, and vice versa). Notably, the four contrast images were also calculated separately for two experimental sessions (i.e., the different- and the same-dimension distractor condition).

In the second-level group analysis, we first identified brain regions that were generally, across the two sessions, involved in a specific condition and then used these as masks for performing the respective test within the two (distractor-type) sessions since we were interested in condition-specific differential responses within the ‘distraction network’. That is, we first submitted the four individual contrast images that combined the same regressors across the two sessions (e.g., distractor present > distractor absent) to one-sample t-tests in order to determine common brain regions activated in one particular condition at a height threshold of *p* < .001 (uncorrected). Next, we used those activated regions as a mask for examining the same condition separately in each experimental session (e.g., distractor present > distractor absent in the different- and, respectively, the same-dimension session) on the group level. Restated, the four individual contrast images within the different- and, respectively, the same-dimension session were taken to the group level and subjected to a one-sample t-test based on the corresponding mask, with family-wise error (FWE) corrected at a cluster-defining voxel-level cut-off of *p* < 0.05 and a minimum cluster size of 5 contiguous voxels.

##### Volume-of-Interest (VOI) analysis

Functional MRI data of the localizer stimuli (checkerboards) at the four positions corresponding to potential target/distractor locations were used to identify localized activation in early visual cortex (see Figure 1C). The first-level GLM model was estimated with four experimental regressors defined by the onset of visual stimulation at each of the four localizer positions, with a duration of 16 s. The hemodynamic response was again modelled by the canonical hemodynamic response function and its first derivative. Six head-movement parameters were included as covariates. Four individual contrast images were calculated by comparing each positional regressor with the other three regressors and then taken to the group level for one-sample t-tests at an extent threshold of p < 0.05 (FWE corrected) with a minimum cluster size of 5 contiguous voxels (Bertleff et al. 2016). The significantly activated clusters thus obtained turned out somewhat different in volume size for the four position localizers. To ensure identical volume sizes for the subsequent VOI analysis, the four localizer VOIs were defined as four spheres, with the centre point of each sphere placed on the peak coordinate defined by the group maximum t value within the respective cluster and with a radius of 9 mm (see Figure 1C, right). The spheres’ radius was determined based on the minimum volume size – consisting of 116 voxels – identified in a group-level analysis of the four localizer positions. In the next step, another set of first-level GLM models were estimated with four experimental regressors representing a *distractor* occurring at one of the localizer positions, separately for two distractor-type sessions. The hemodynamic response related to neural activity in the four distractor regressors was modelled by the canonical hemodynamic response function and its first derivative, with six head-movement parameters considered covariates in the model. Analogous GLM models were developed with four regressors representing a *target* appearing at one of the four positions, separately for two distractor-type sessions. Percent signal change (beta values) of experimental regressors were extracted within the corresponding localizer VOIs for further examination.

## Results Behavioural Results

The first trial in each block was excluded from analysis, as were response-error trials in the response-time (RT) analysis.

The error rate was overall higher in the same-dimension than in the different-dimension session (14.47% vs. 12.66%); and, compared to the distractor-absent baseline (10.7%), more errors were made on trials in which a distractor was present (in the rare region: 16.3%; in the frequent region, 15.7%). Further, the increased error rates caused by distractor presence were more marked with same-than with different-dimension distractors (see Figure 2a). This effect pattern was confirmed by a repeated-measures ANOVA with the factors Distractor (absent, present in the frequent region, present in the rare region) and Distractor Type (different-dimension distractor, same-dimension distractor), which revealed all effects to be significant: Distractor, *F*(2, 62) = 32.49, *p* < .001, .109; Distractor Type, *F*(1, 31) = 6.41, *p* = .017, .034; interaction, *F*(2, 62) = 11.24, *p* < .001, .032.

**Figure 2.**
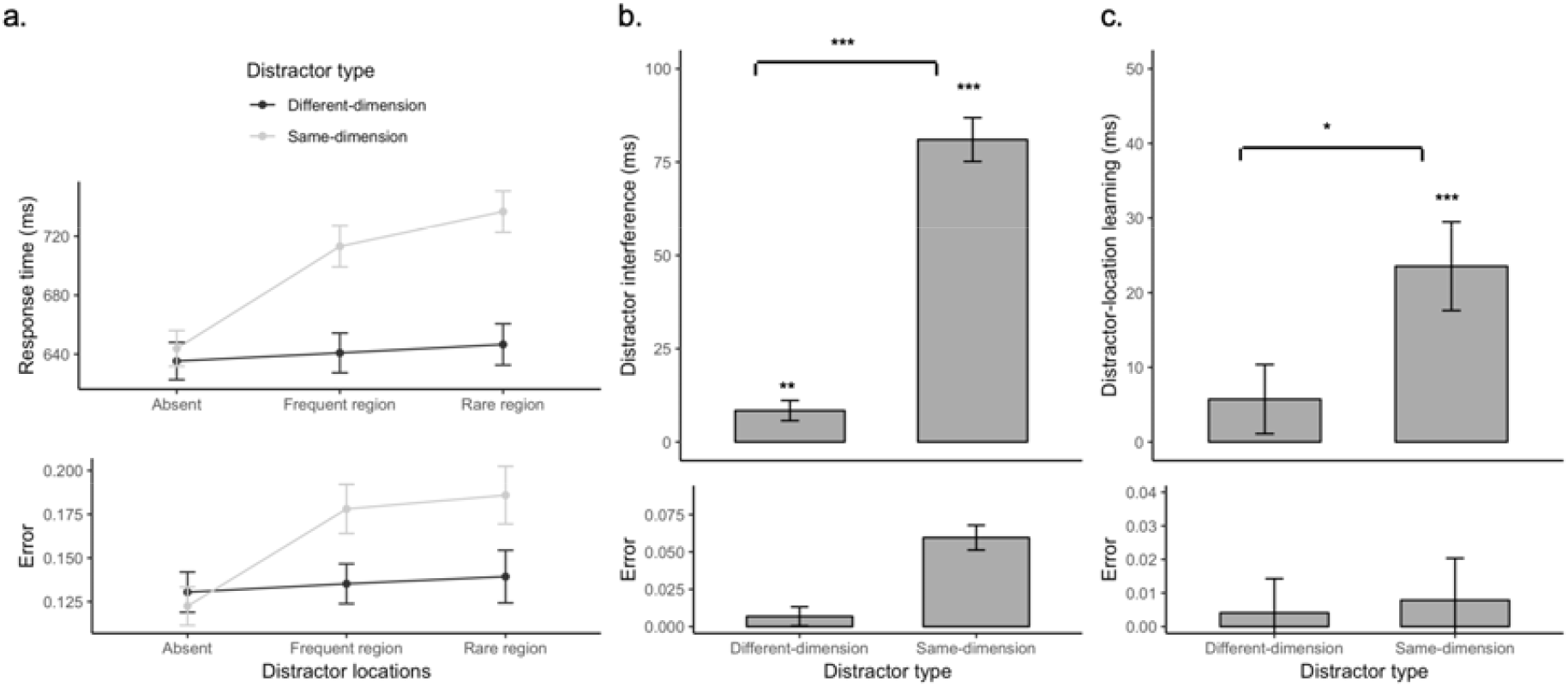
Response times (RTs; upper panels) and error rates (lower panels) for the two distractor types. (a) Averaged RTs and error rates in the three distractor conditions, separately for the different- and same-dimension distractor sessions. (b) Distractor-interference effect, calculated as the difference between distractor-present and -absent trials, separately for the different- and same-dimension distractors. (c) Distractor-location learning effect, calculated as the difference between trials with a distractor presented in the rare vs. frequent region, separately for the different- and same-dimension distractors. Error bars depict 95% confidence intervals. * denotes p < .05, ** p < .01, *** p < .001.

This (interactive) effect pattern was mirrored in the RT results (Figure 2a), effectively ruling out differential speed-accuracy trade-offs. An analogous ANOVA of the mean RTs again revealed all effects to be significant: Distractor, *F*(2, 62) =122.6, *p* < .001, .082; Distractor Type, *F*(1, 31) = 59.9, *p* < .001, .131; interaction, *F*(2, 62) = 59.4, *p* < .001, .054. Response speed was overall slower with same-than with different-dimension distractors, and the presence of a distractor slowed RTs to the target (relative to the distractor-absent baseline). This slowing was more marked in the same-than in the different-dimension distractor condition; as depicted in Figure 2b, the interference effect was only some 8 ms with different-dimension distractors, *t*(31) = 3.15, *p* = .004, but ten times as high (81 ms) with same-dimension distractors, *t*(31) = 14.0, *p* < .001. This differential interference effect was significant (*t*(31) = -12.2, *p* < .001). Thus, even though the two types of distractor were balanced in terms of bottom-up saliency, same-dimension distractors caused considerably more RT interference than different-dimension distractors – in line with previous findings (e.g., Sauter et al. 2018, 2019; Liesefeld and Müller 2020).

To quantify the effect of distractor-location learning, we calculated the RT difference between trials with a distractor presented in the rare region versus trials with a distractor presented in the frequent region. As depicted in Figure 2c, when a distractor appeared in the frequent region, RTs to the target were generally faster than with a distractor appearing in the rare region. Importantly, this difference was greater with same-dimension distractors, evidenced by a significant distractor-location effect in the same-dimension session (24-ms benefit, *t*(31) = 4.03, *p* < .001), but not in the different-dimension condition (6-ms benefit, *t*(31) = 1.26, *p* = .218). In any case, the larger (frequent-vs. rare-region) RT benefits obtained with same-than with different-dimension distractors (*t*(31) = -2.04, *p* = .05) closely replicate our previous findings (e.g., Sauter et al. 2018, 2019; Liesefeld and Müller 2020).

Of note, even though the target occurred with equal likelihood in both distractor regions, targets appearing at a location in the frequent region were responded to slower than targets in the rare region, the RT costs amounting to some 9 ms (combined across distractor-present and -absent trials) with different-dimension distractors (*t*(31) = 2.61, *p* = .014) and to 18 ms with same-dimension distractors (*t*(31) = 4.31, p < .001). Although the RT cost was double the size in the same-versus the different-dimension condition, the difference was non-significant (*t*(31) = -1.23, *p* = .228). Thus, while statistical learning of distractor locations reduced the interference caused by distractors in the frequent region, this was associated with a cost: slowed processing of targets appearing in the frequent (distractor) region. Consistent with our previous behavioural studies, this cost effect was more marked, at least numerically, with same-dimension distractors. [In previous studies, there was either no cost with different-dimension distractors (e.g., Liesefeld and Müller 2020), or there was some cost initially, which, however, disappeared over extended task practice (Zhang et al. 2019).]

### VOI Results

Based on human probabilistic cytoarchitectonic maps within the Anatomy Toolbox (Eickhoff et al. 2005), the group peak coordinates of the maximum *t*-values associated with each of the four flickering checkerboard localizers – that is, potential target/distractor positions – were localized to early visual cortex (V1 – V4; Figure 1C, right).

We first examined changes in the beta values representing activation at the specific localizer positions (VOIs) when the distractor appeared at a location in the frequent and, respectively, the rare region, for the two distractor types. To start with, we submitted the beta values to a three-way ANOVA with the within-subject factors Distractor Region (frequent region, rare region) and Distractor Type (different-, same-dimension distractor) and the between-subject factor Frequent Hemisphere (Group 1 with different-dimension distractors frequently appearing in the left region, i.e., the right VOIs, and same-dimension distractors frequently appearing in the right region, i.e., the left VOIs; Group 2 with the reversed frequent hemisphere relative to Group 1 for two distractor types). As the effect of the distractor-frequency manipulation did not differ between the two groups (non-significant main effect of Frequent Hemisphere, non-significant Frequent Hemisphere × Distractor Region or, respectively, Frequent Hemisphere × Distractor Type interactions, all *p*s > .07), we collapsed the beta values across the different assignments of the frequent distractor regions.

Figure 3 depicts the resulting beta values for distractor locations in the frequent and, respectively, rare distractor regions, separately for each distractor type. By visual inspection, and as confirmed by repeated-measures ANOVA of Distractor Type and Distractor Region, the beta values were overall lower for distractors appearing in the frequent versus the rare region (significant main effect of Distractor Region, *F*(1, 25) = 7.57, *p* = .01). This pattern is consistent with the idea that statistical learning of distractor locations is associated with stronger signal suppression in early visual areas coding frequent versus rare distractor locations. However, in contrast to the RT results, the beta values turned out little influenced by the factor Distractor Type (main effect, *F*(1, 25) = 1.24, *p* = .28); in particular, the effect of (frequent, rare) distractor region did not appear to be reduced in the different-, as compared to the same-, dimension distractor condition (non-significant Distractor Type × Distractor Region interaction, *F*(1, 25) = 0.09, *p* = .76). However, as a weaker effect was expected from the RT pattern, we conducted paired t-tests comparing the beta values between the frequent and rare distractor regions, separately for the two distractor types. These revealed the difference to be significant for the same-dimension condition (rare vs. frequent region: 0.06 vs. -0.27, *t*(25) = 2.45, *p* = .022), but not for the different-dimension condition (0.35 vs. 0.09, *t*(25) = 1.57, *p* = .13). Thus, while early visual-cortex activation was generally reduced when a distractor occurred in the frequent (vs. the rare) region, this effect was statistically robust (i.e., consistent across participants) only with same-dimension distractors, but not with different-dimension distractors.

**Figure 3.**
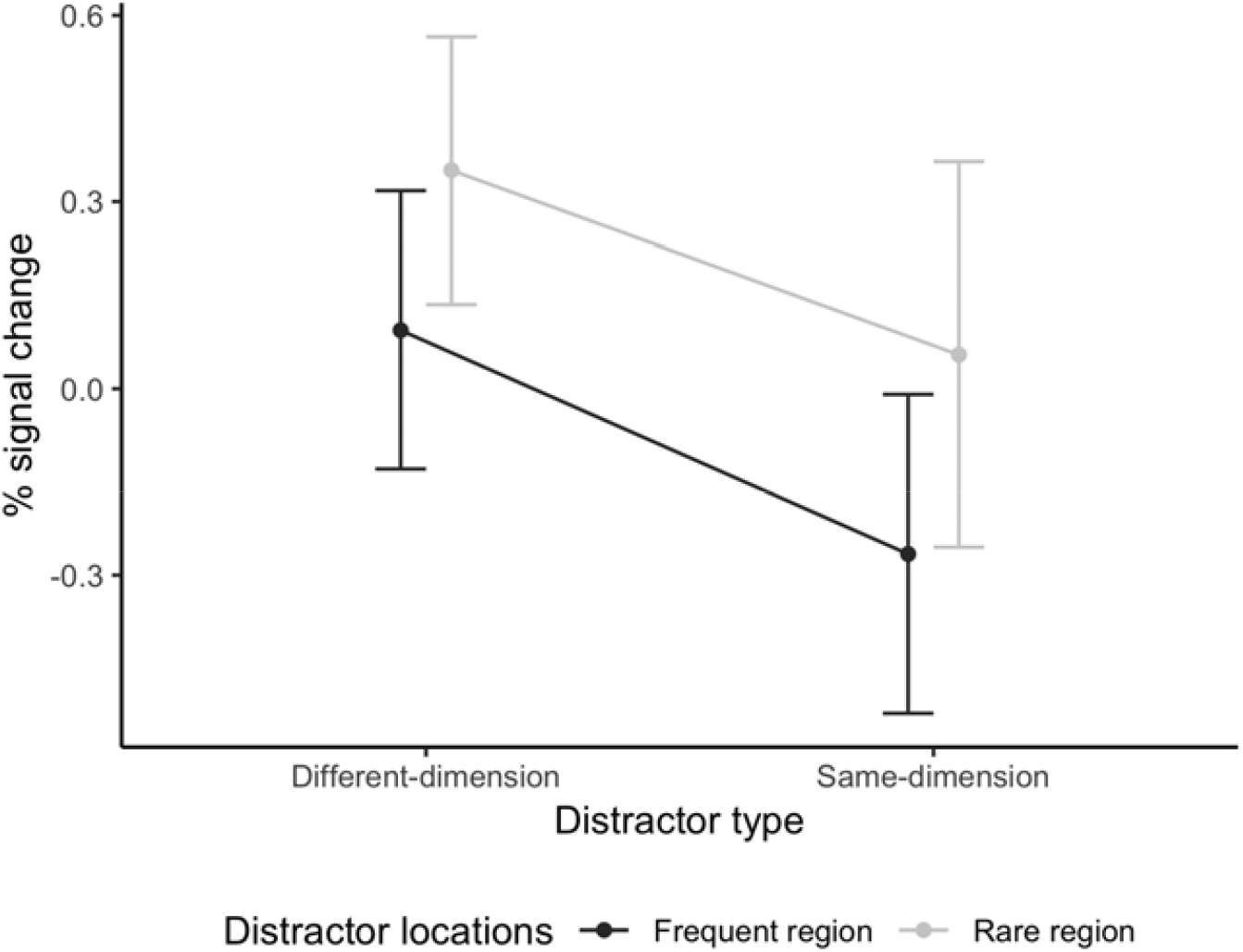
Mean percent signal change (beta values) representing early visual activation by singleton distractors appearing in the frequent vs. the rare distractor region, separately for the different- and same-dimension distractor types. Error bars depict 95% confidence intervals.

Given this, we further examined whether the early visual-cortex modulations play a role in generating the behavioural effects. To this end, we analysed the relationships between the beta values and the RT interference caused by distractors occurring in the frequent and, respectively, the rare region, for each of the two distractor-type conditions. The correlations are illustrated in Figure 4. As can be seen, the beta values were predictive of RT-interference magnitude only in the same-dimension condition (Frequent region: *r* = .517, *p* = .007; Rare region: *r* = .466, *p* = .016), but not the different-dimension condition (Frequent region: *r* = .094, *p* = .646; Rare region: *r* = -.052, *p* = .800). This pattern points to a critical role of the early visual signal modulations for behavioural distractor interference only with same-dimension distractors; in contrast, some other, or additional, distractor-handling mechanism must come into play with different-dimension distractors (see whole-brain results below).

**Figure 4.**
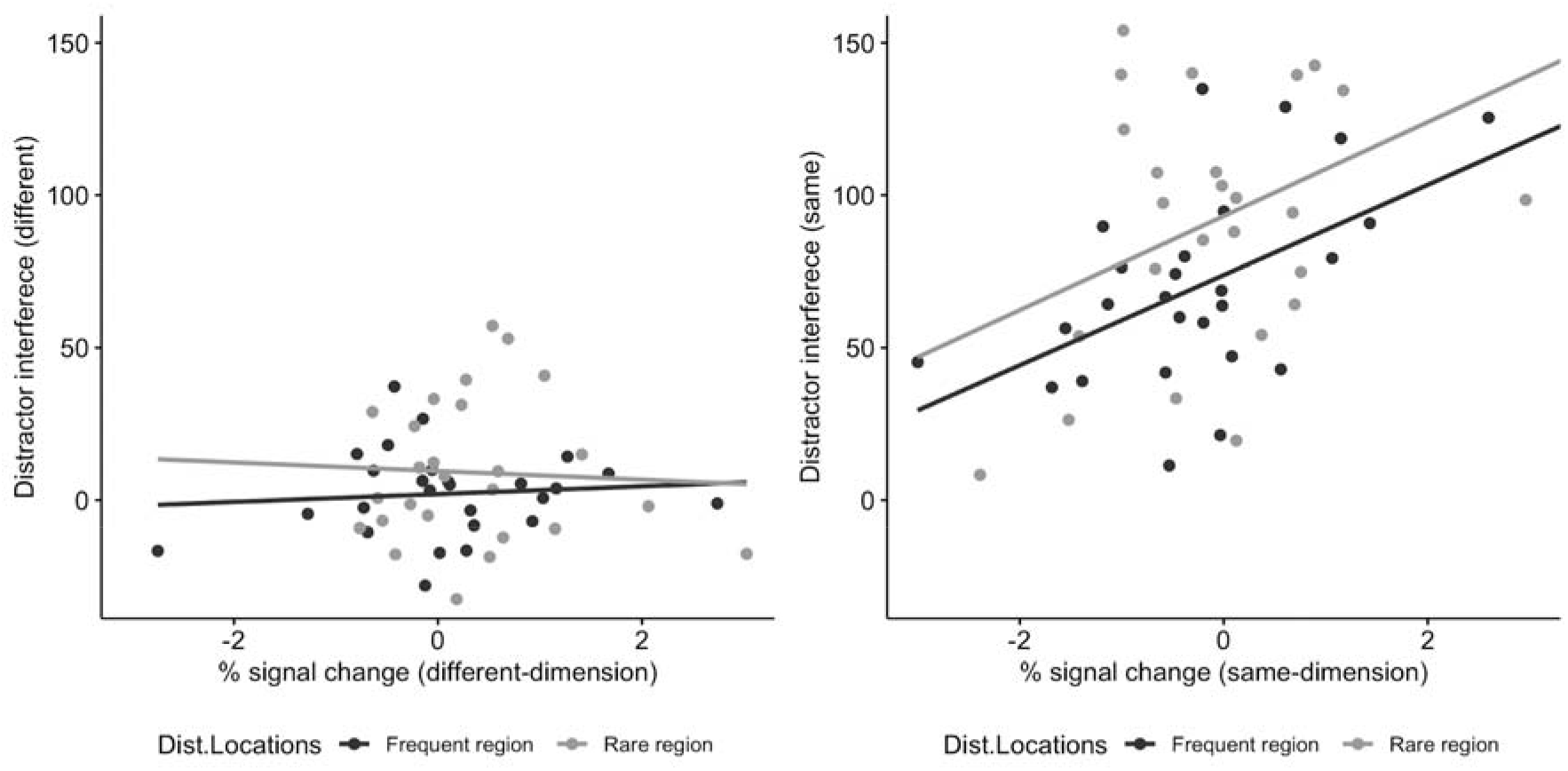
Correlation between behavioural distractor interference effect (RTs) in the frequent region and the rare region with the respective percent signal changes for distractors in the frequent and rare region, separately for the different- (left panel) and the same-dimension distractor types (right panel).

Of note, the beta values were not only reduced when a distractor occurred in the frequent (vs. the rare) region (see above), but also when a target appeared there (significant main effect of Target Location in frequent vs. rare region: *F*(1, 25) = 6.90, *p* = .015). Although the beta values were numerically more negative overall in the same-dimension condition, the main effect of Distractor Type was non-significant (*F*(1, 25) = 0.51, *p* = .48). Finally, the reduction was comparable between the two distractor-type conditions (Target-Location × Distractor-Type interaction: *F*(1, 25) = 0.008, *p* = .93), even though it tended to be more robust in the same-dimension (rare vs. frequent region: 0.09 vs. -0.22, *t*(25) = -2.42, *p* = .023) than in the different-dimension distractor condition (0.33 vs. 0.01, *t*(25) = -1.85, *p* = .077). This pattern is similar to the distractor-location effects (see above), and so likely reflecting the same mechanisms underlying statistical distractor-location learning.

### Whole-brain Results

Whole-brain results showed that the presence of a singleton distractor defined in a different dimension (namely colour) to the target (orientation) invoked a BOLD response in left fusiform gyrus (FWE corrected, *p* < .05, see Figure 5 and Table 1). Furthermore, compared to a (colour) distractor appearing in the frequent region, a distractor in the rare region induced more robust activation in the right superior parietal lobule (Brodmann area, BA 7), left fusiform gyrus, as well as large parts of the occipital cortex (FWE corrected, *p* < .05, see Table 1, Figure 5). In contrast to the different-dimension distractor, the presence of a distractor defined in the same dimension as the target was associated with more robust activation in the right superior parietal lobule (BA 7) as well as the left superior parietal lobule extending to left middle occipital regions (FWE corrected, *p* < .05, see Table 1, Figure 5). Critically, however, no significant clusters were found when comparing (same-dimension) distractors in the rare region versus the frequent region. This pattern suggests that distractor handling in general and statistical distractor-location learning in particular operate more in early visual cortical areas with same-dimension (orientation-defined) distractors (see VOI results above), whereas some higher-level mechanism comes into play with different-dimension (colour-defined) distractors.

**Table 1.** List of activations associated with contrasts defined by (A) Distractor present > absent, (B) Distractor in the rare region > frequent region, (C) Distractor in the frequent region > rare region, separately for the different- and same-dimension distractor types.

| *Contrast* | *Side* | *Region* | *Cluster size* | *Cluster peak coordinates* | *T value* |
| --- | --- | --- | --- | --- | --- |
| *Different-dimension session* |  |  |  |  |  |
| (A) Distractor present > absent | L | Fusiform gyrus | 7 | -33, -57, -12 | 4.70 |
| (B) Distractor in the rare region > frequent region | L | Superior occipital lobule | 16 | -21, -63, 30 | 4.92 |
|  | R | Middle occipital gyrus | 26 | 30, -69, 24 | 4.67 |
|  | R | Inferior occipital lobe | 6 | 36, -66, -12 | 4.61 |
|  | L | Fusiform gyrus | 30 | -36, -60, -12 | 4.63 |
|  | R | Superior parietal lobule (BA 7) | 17 | 30, -57, 51 | 4.59 |
| (C) Distractor in the frequent region > rare region | No significant brain activation |  |  |  |  |
| *Same-dimension session* |  |  |  |  |  |
| (A) Distractor present > absent | L | Superior occipital lobule | 68 | -21, -60, 51 | 5.20 |
|  | L | Superior occipital lobule | 10 | -24, -69, 27 | 5.03 |
|  | R | Superior parietal lobule (BA 7) | 26 | 27, -63, 45 | 4.02 |
| (B) Distractor in the rare region > Frequent region | No significant brain activation |  |  |  |  |
| (C) Distractor in the frequent region > Rare region | No significant brain activation |  |  |  |  |

Coordinates (x, y, z) were defined in MNI space. Activations were all significant at $p < 0.05$ (FWE corrected), at the cluster level (based on $p < 0.001$ , at the voxel level).

**Figure 5.**
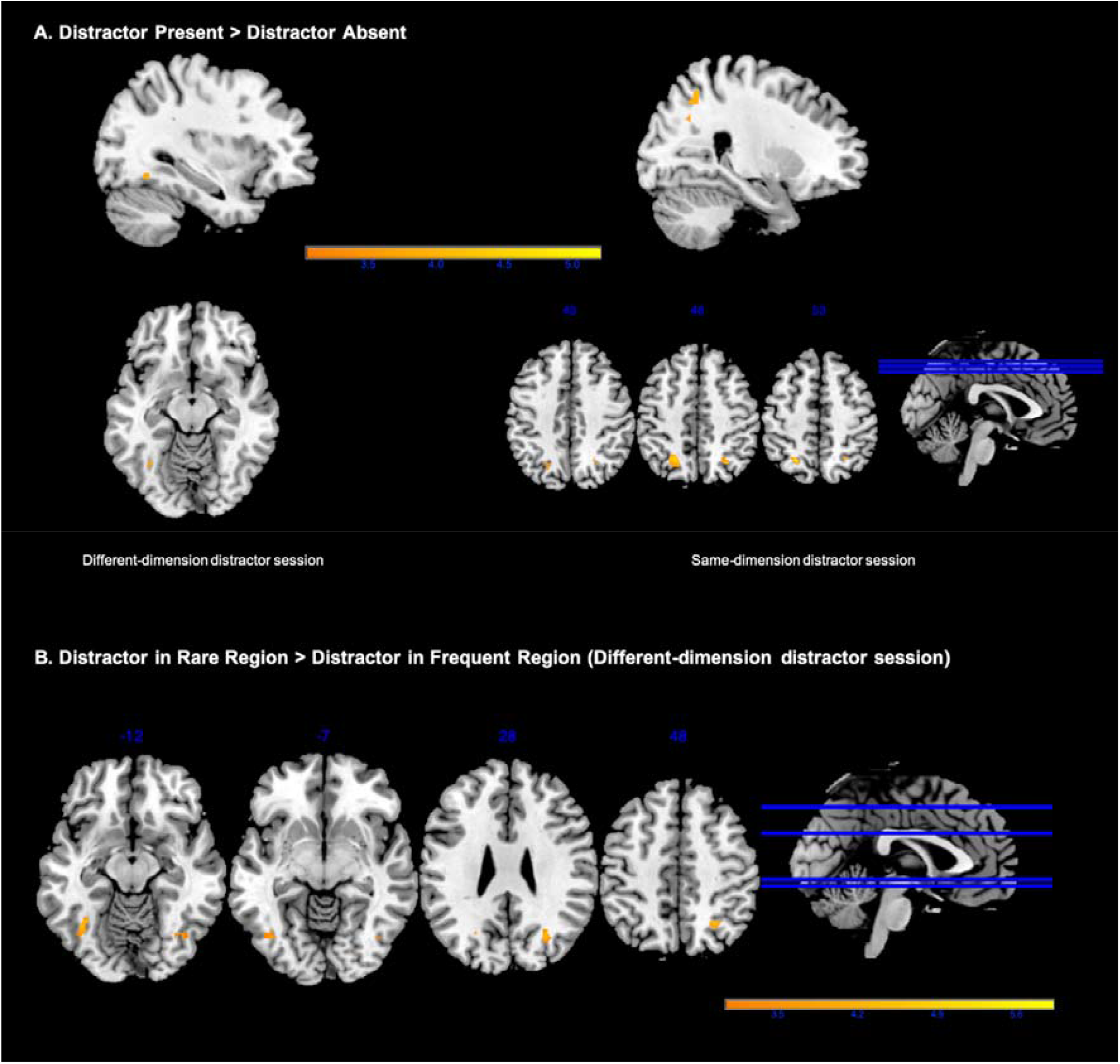
**A**. Whole-brain activation patterns coloured in yellow reflect invoked BOLD signals driven mainly by the presence of salient distractors defined in a different dimension (namely, colour) to the search-relevant target (left, different-dimension session), and, respectively, the presence of distractors defined in the same dimension (namely, orientation) as the target (right, same-dimension session) at p < .05, family-wise error (FWE) corrected at the cluster level. **B**. Whole-brain activation patterns coloured in yellow depict increased BOLD signals when different-dimension (i.e., colour-defined) distractors appeared in the rare as compared to the frequent region, at p < .05, FWE corrected at the cluster level.

## Discussion

Combining fMRI with a statistical distractor-location learning paradigm, we manipulated whether the singleton distractor was defined within the same dimension (orientation) or a different dimension (colour) relative to the target. The behavioural results replicated previous findings: interference by a salient distractor was reduced when it appeared within the frequent, versus the rare, distractor region – evidencing adaptation of attentional guidance to the biased distractor distribution. Further, despite being equally (if not more) salient, different-dimension distractors produced substantially less interference than same-dimension distractors, associated with a less marked frequent versus rare distractor-region effect. These behavioural effects were, to some extent, reflected in the fMRI results. BOLD signals in the early visual cortex were reduced for distractors occurring in the frequent versus rare region. While the reduction was numerically similar in the two distractor conditions, it was more robust with same-dimension distractors, and crucially, behavioural interference correlated with distractor-evoked VOI activity exclusively for this type of distractor. A similar activity pattern was evident when (spatially unbiased) targets appeared in the frequent versus rare distractor region, mirroring a similar effect in the RTs. Whole-brain analysis revealed the involvement of parietal parts of the fronto-parietal attention network in distractor handling. Importantly, though, in the different- (but not same-) dimension distractor condition, fusiform gyrus was activated more when a distractor was present versus absent and more with a distractor occurring in the rare versus frequent region. This suggests that distractors defined in a different dimension to the target (namely, colour) are, in crucial respects, handled differently by the brain to same-dimension distractors.

The behavioural signature of statistical distractor-location learning has been well documented recently: RT interference is reduced for distractors occurring at frequent versus rare locations, and this is associated with reduced capture of the first saccade by distractors at frequent locations (Di Caro et al. 2019; Wang et al. 2019a; Sauter et al. 2020). Together with an ERP component interpreted in terms of distractor suppression (Wang et al. 2019b), this has been taken as evidence that observers learn to down-modulate the attentional priority signals (Itti and Koch 2001; Fecteau and Munoz 2006; Wolfe and Gray 2007; Kok et al. 2012; Aitken et al. 2020) generated by distractors at frequent locations, thus reducing their potential to capture attention and cause interference. In line with this, we found that early visual-cortex signalling was reduced for distractors occurring in the frequent versus rare region. Assuming that the attentional priority map is situated at some superordinate level in the visual system – such as the pulvinar thalamus, which is thought to integrate saliency signals from, e.g., LIP, FEF (Bundesen et al. 2005) – reduced distractor signalling in early visual cortex might reflect learnt top-down inhibition of feature coding in early visual areas. The fact that this is observed generally (with both types of distractor) is consistent with Won et al. (2020), who found reduced visual-cortex signalling when different-dimension distractors (i.e., colour singletons that varied in the specific colour feature) occurred with 80%, but not 25%, frequency anywhere in the search display. In contrast, the reason why Bertleff et al. (2016) did not find evidence of down-modulated distractor signalling (when comparing blocks with 100% vs. 0% distractor presence) in early visual areas may be that they varied the spatial-attentional setting (focused vs. distributed) for the *target* (rather than the *distractor*), along with the use of different-dimension (colour) distractors.

Neurally, input coding in early visual cortex is thought to constitute the first computational stage of salience processing: the generation of local feature-contrast, or ‘saliency’, signals (Knierim and van Essen 1992; Nothdurft 2000; Li 2002) within the various feature dimensions, which are subsequently integrated across dimensions into an ‘overall-saliency’ map determining the priorities for the allocation of attention. Stimuli that contrast more strongly with their surroundings (i.e., are more bottom-up salient) generate higher peaks on the priority map and have a higher likelihood of summoning attention (Treue 2003; Töllner et al. 2011; Kamkar et al. 2018). Accordingly, if distractors are more salient than targets, they are more likely to capture attention inadvertently. Thus, our finding of reduced distractor signals in the early visual cortex (especially at frequent locations) likely indicates a general down-modulation of feature-contrast signals, broadly consistent with Gaspelin and Luck’s (2019) ‘signal-suppression’ hypothesis.

Of note, if anything, our colour distractors were more salient than our orientation distractors (see Method), and so, on a purely bottom-up account, they should *not* have produced less interference than the orientation distractors. However, the fact that the colour distractors produced substantially less (rather than more) behavioural interference, coupled with a less marked frequent versus rare distractor-region RT effect and the absence of a correlation of early visual-cortical BOLD activity with the magnitude of RT interference, suggests that distractor-signal suppression, and particularly enhanced suppression in the frequent versus rare region, involved some other or additional mechanism with different-dimension distractors.

According to the Dimension-Weighting Account (Found and Müller 1996; Müller et al. 2003, 2009; Liesefeld and Müller 2020), such a mechanism is provided by dimension-based signal suppression (also referred to as ‘second-order feature suppression’ by Gaspelin and Luck 2018; Won et al. 2019). That is, with distractors defined in a different dimension to the target (here: colour distractors, orientation targets), suppression might operate at the level of the distractor dimension, selectively down-modulating the contribution of colour signals to (supra-dimensional) priority computation without impacting the contribution of orientation signals. This strategy is unavailable with same-dimension distractors. Consistent with a filtering stage specific to the distractor dimension, the whole-brain analysis revealed that the left fusiform gyrus was generally involved in dealing with colour-defined different-dimension distractors (whereas it was not activated by orientation-defined same-dimension distractors). Previous neuropsychological, electrophysiological, and neuroimaging studies have revealed the (left) fusiform gyrus to play a role in colour processing (Allison et al. 1993; Chao and Martin 1999; Pollmann et al. 2000; Simmons et al. 2007). Of note, Simmons et al. (2007) considered the left fusiform gyrus to be “a high-level colour perception region” that is activated not only during colour perception (responding more strongly to colour than to greyscale stimuli), but also during the top-down-controlled retrieval of conceptual colour knowledge (i.e., during verifying whether a named colour is true of a named object). In the present study, the left fusiform gyrus was generally activated by colour distractors, compared to when no distractors were present in the display. This pattern is consistent with the fusiform gyrus playing a role in colour-based stimulus filtering via reducing the weight of colour-based feature-contrast signals in attentional priority computation. Previous studies have shown that colour-based distractor filtering can operate quite effectively across all display locations (e.g., Müller et al. 2009; Won et al. 2019), so tuning the filter to a region where colour distractors occur frequently might yield little extra benefits. Spatially uniform filtering could explain why the correlation between distractor-generated BOLD activity in early visual areas and behavioural (RT) distractor interference was effectively abolished for colour-defined distractors (while it was robust for orientation-defined distractors). Additionally, the *dimensional* filter might be modified by statistical distractor-location learning, up-modulating the suppression weights for colour signals in the frequent, relative to the rare, distractor region. Consistent with this, the fusiform gyrus was activated less strongly by colour distractors in the frequent versus the rare region. – Given this general sketch of learnt distractor suppression, at least two questions arise: 1) How does the adaptation, in the early visual cortex, to the spatial distractor distribution come about? 2) Why was the beta-value gradient between the frequent- and rare-distractors-region VOIs not significantly reduced for different-compared to same-dimension distractors?

Concerning the first question, the reduced response to distractors in the frequent versus the rare region might reflect a form of low-level ‘habituation’ (e.g., Turatto et al. 2018). Of note, though, VOI activity was reduced not only to distractor signals in the frequent region but also to target signals (despite targets occurring with equal frequency in both regions). Behavioural work has demonstrated *facilitation* of locations at which *targets* frequently appear, analogous to *inhibition* of positions where *distractors* frequently occur (Ferrante et al. 2018) – suggesting that behavioural facilitation (target-location learning) is the flipside of inhibition (distractor-location learning). Thus, if inhibition involves a top-down-mediated reduction of neural responsivity in the early visual cortex, owing to the status of ‘distractors’ as task-irrelevant, to-be-rejected items, one would expect facilitation to be associated with *higher* beta values for targets at frequent versus rare *target* locations. This expectation is at variance with habituation accounts predicting the beta values to be *lower* (as for distractors at frequent versus rare *distractor* locations). To our knowledge, these contrasting predictions have not yet been tested for statistical *target*-location learning. However, assuming that *distractor*-location inhibition is top-down mediated (tied to the status of distractors as ‘distractors’), the fact that target signals, too, were reduced in the early visual cortex would argue in favour of the inhibition at the lower level originating from some higher level. One likely source is the priority map, that is, inhibition of salient distractors that captured attention at the priority-map level feeds back to and adapts (‘habituates’) neuronal responsivity in feature-coding areas. Consistent with the notion of the priority map being an inherently ‘feature-blind’ representation, this feeding-back of inhibition appears to be feature-unspecific: it impacts not only the coding of the distractor feature but also that of the target feature, even if the latter belongs to a different dimension. Of note, though, the feedback tended to be generally weaker in the different (vs. the same-) dimension condition, as reflected by the beta values being numerically more positive for VOIs in both the frequent and rare distractor regions (this pattern was seen both with a distractor and a target appearing in a given VOI). Weaker feedback is also consistent with a reduced target-location effect in the different- (vs. the same-) dimension condition.

A second question concerns why the beta-value gradient between the frequent- and rare-distractor-region VOIs was not noticeably reduced for different-compared to same-dimension distractors in the present study, given that different-dimension distractors permitted efficient, dimension-based filtering of distractor signals. One possibility is that the gradient is learnt early on during practice (e.g., already in the first trial block, because distractors in the frequent region capture attention more often than distractors in the rare region) and then persists. In contrast, the strategy of dimension-based filtering would be ‘discovered’ only later on, once the early-level gradient has been established (Zhang et al. 2019). That is, capture prevention by dimension-based filtering does not lead to unlearning of the initially acquired gradient. Consistent with this are indications that statistical distractor-learning effects are pretty resistant to unlearning (Turatto et al. 2018). Alternatively, even after a different-dimension distractor seizes to capture attention (due to efficient dimension-based filtering), the presence of a distractor may still be registered and responded to with top-down inhibitory feedback, reinforcing the gradient at the lower level. That is, the gradient reflects distractor frequency in the two regions, independently of whether or not the distractor is potent enough to capture attention. In other words, the low-level gradient represents the basic distractor-region ‘prior’.

The whole-brain analysis also revealed the right SPL to be more strongly activated by different-dimension distractors appearing in the rare versus the frequent region (an effect not seen with same-dimension distractors). The right SPL, which has long been considered critical for visual-spatial attentional control (Shapiro et al. 2002; Thakral and Slotnick 2009), is engaged not only in shifts of spatial attention (Corbetta et al. 1995; Behrmann et al. 2004) but also in shifting attention between separable dimensions of the input (Yantis and Serences 2003). The more robust SPL activation by different-dimension distractors in the rare region might reflect a higher incidence of attentional capture by such distractors, which may require combined dimensional and spatial shifting of attention to a target defined in a different dimension. Dimensional shifting would not be required with same-dimension distractors, which might explain why no distractor-region-specific SPL activation was seen in the same-dimension distractor condition.

In summary, the current results show that statistical learning of distractor locations involves (acquired) suppression down to the level of the early visual cortex. Besides, with different-dimension (colour) distractors, higher-level, dimension-specific filtering mechanisms can come into play. Colour-based filtering, involving the right fusiform gyrus and SPL, substantially reduces the interference caused by colour distractors, whether they occur in the frequent or rare region. A dimension-based filtering strategy does not seem to be available with distractors defined in the same dimension as the target (orientation), in which case interference reduction relies solely on spatially tuned lower-level signal suppression.

